# Demographic Consequences of Damage Dynamics in Single-Cell Ageing

**DOI:** 10.1101/2023.05.23.538602

**Authors:** Murat Tuğrul, Ulrich K Steiner

## Abstract

Ageing is driven by damage accumulation leading to a decline in function over time. In single-cell systems, in addition to this damage accumulation within individuals, asymmetric damage partitioning at cell division can play a crucial role in shaping demographic ageing patterns. Despite empirical single-cell studies providing quantitative data at the molecular and demographic level, a comprehensive theory of how cellular damage production and asymmetric partitioning propagate and influence demographic patterns is still lacking. Here, we present a generic and flexible damage model using a stochastic differential equation approach that incorporates stochastic damage accumulation and asymmetric damage partitioning at cell divisions. We formulate an analytical approximation linking cellular and damage parameters to demographic ageing patterns. Interestingly, the lifespan of cells follows an inverse-gaussian distribution whose underlying properties derive from cellular and damage parameters. We demonstrate how stochasticity (noise) in damage production, asymmetry in damage partitioning, and division frequency shape lifespan distribution. Confronting the model to various empirical *E*.*coli* data reveals non-exponential scaling in mortality rates, a scaling that cannot be captured by classical Gompertz-Makeham models. Our findings provide a deep understanding of how fundamental processes contribute to cellular damage dynamics and generate demographic patterns. Our damage model’s generic nature offers a valuable framework for investigating ageing in diverse biological systems.

**Significance:** Asymmetries and randomness in cellular events play important roles in establishing the diversity at evolutionary and demographic scales. Looking at single-cells, ageing processes are influenced by stochastic damage accumulation and asymmetric damage partitioning at cell divisions. Utilising stochastic differential equations, we develop a cellular damage model that encapsulate both noisy damage accumulation within cells and asymmetric damage partitioning among cells when they divide. In doing so we bridge molecular stochastic processes, demographic fates of individuals, and population level demographic distributions. Our model aligns with empirical data from *E*.*coli* single-cell studies and advances our understanding compared to traditional demographic models. The generic and adaptable nature of our model paves the way for broader applications in ageing research across biological systems, highlighting the influences of stochasticity and asymmetry on cellular biology.

## INTRODUCTION

Ageing disrupts cellular integrity and leads to functional decline in cells and individuals. Despite its wide-ranging effects from medicine to evolution, our mechanistic understanding of ageing linking from the molecular levels to the demographic levels is still limited [López-Otín et al. 2013]. At *micro* scales, ageing is considered to be driven by damage accumulation interfering cellular processes. Damage sources include DNA oxidation, protein aggregation, mitochondrial dysfunction and mutations [Gladyshev 2016, Schumacher et al. 2021]. At *macro* scales, demographic ageing patterns are surprisingly diverse across species, typically with high variation in lifespans [Jones et al. 2014]. Molecular studies of ageing mostly focused on genetics and environment to understand this variation [López-Otín et al. 2013] but there remains a neglected stochastic component in demographic variation [Steinsaltz et al. 2020]. This stochastic heterogeneity in ageing is illustrated by genetically identical twin and inbred animal studies [Finch and Kirk- wood 2000], as well as isogenic individuals in highly controlled environments [Wang *et al*. 2010]. Inherent stochasticity in cellular processes such as noisy gene expression [Elowitz *et al*. 2002, Raj and Oudenaarden 2008, Patange *et al*. 2018] and asymmetric cell divisions [Huh and Paulsson 2011a,b, Shi *et al*. 2020] might be the ultimate sources of this demographic heterogeneity. High-throughput single-cell studies and mathematical models are crucial to decipher the relationship between stochastic cellular damage dynamics and demographic variation in ageing.

In their seminal work, Stewart *et al*. [2005] showed that *E*.*coli*, a morphologically symmetric dividing microbe, is also susceptible to reproductive ageing. This illustrates that ageing goes beyond morphologically asymmetric and sexually reproducing organisms and provides us with a model organism to study individual heterogeneity in ageing with available molecular, genetic, and imaging techniques. Bacterial ageing (for review, see [Steiner 2021]) has been best explored by experimental studies on cellular growth and division in ageing under external stress [Wang *et al*. 2010, Rang *et al*. 2011, 2012, Łapińska *et al*. 2019] and by molecular factors such as chaperones associated with protein aggregation [Winkler et al. 2010, Proenca et al. 2019] Theoretical models of bacterial ageing focused on the optimality of asymmetric damage partitioning for maximising fitness [Watve et al. 2006, Evans and Steinsaltz 2007]; distinct reproductive growth states [Chao 2010, Blitvić and Fernandez 2020], and damage repair [Clegg et al. 2014]. Longer lasting single-cell experiments added on cell survival and mortality insights [Wang et al. 2010, Jouvet et al. 2018, Robert et al. 2018, Yang et al. 2019, Steiner et al. 2019]. These studies confirmed chronological ageing in bacteria and showed high variations in lifespans deviating from random mortality processes. However, how stochastic damage dynamics leads to individual differences in demographic ageing patterns remains elusive.

Existing mathematical models for bacterial ageing mostly rested within an equilibrium assumption and did not exploit underlying dynamical aspects influencing ageing and demography patterns. In demography, lifespan-related data, e.g. mortality risk chances with age, have been traditionally interpreted with phenomenological models such as the Gompertz-Makeham model [Makeham 1860]. These models not only fail to capture any non-exponential mortality rates but usually do not connect molecular damage mechanisms and demographic patterns. Anderson [2000] and Weitz and Fraser [2001] independently developed stochastic models of vitality using a stochastic differential equation (SDE) approach which explained some common ageing patterns such as mortality plateaus observed in various organisms [Jones et al. 2014]. To our knowledge, a similar SDE modelling framework has not been adopted for single-cell or bacterial ageing, or remained minimal in applications [Yang *et al*. 2023]. Extensions of such SDE models require dealing with asymmetric damage partitioning at cell divisions, i.e. sudden jumps in accumulated intracellular damage.

Here we develop a general framework by adapting jump-diffusion type SDEs for the damage dynamics along mother bacterial cells. This modelled damage dynamics determines the lifespan-related demographic characteristics. We then explore how demographic outcomes are shaped by cellular damage characteristics, i.e. initial damage levels, damage production rate and noise, and damage partitioning asymmetry, as well as cellular division rate and fluctuations. Lastly, we use various empirical dataset for *E*.*coli* with mortality events [Wang et al. 2010, Robert et al. 2018, Yang et al. 2019, Steiner et al. 2019] to demonstrate how such an extended jump-diffusion type SDE model can help us interpret bacterial ageing.

## MODEL

We build a modelling framework based on the assumption that single-cells deteriorate over time due to the accumulation and transmission of *damage*. This damage causes cell death (mortality) upon reaching a critical value. We consider a damage density measure, i.e. number of damage bodies per cell volume, and thereby integrate cellular growth, dilution and repair processes. Here, we focus on modelling the cellular damage dynamics along mother cell lineages (**Fig. 1-a**) which receive the older cell wall (old pole) at cell divisions and typically receive more damage than the daughter cells [Chao *etal*. 2016]. Our model considers two important processes: cellular damage production and partitioning. The damage production encapsulates a deterministic net rate and a stochastic noise for random fluctuations. The damage

**FIG. 1.**
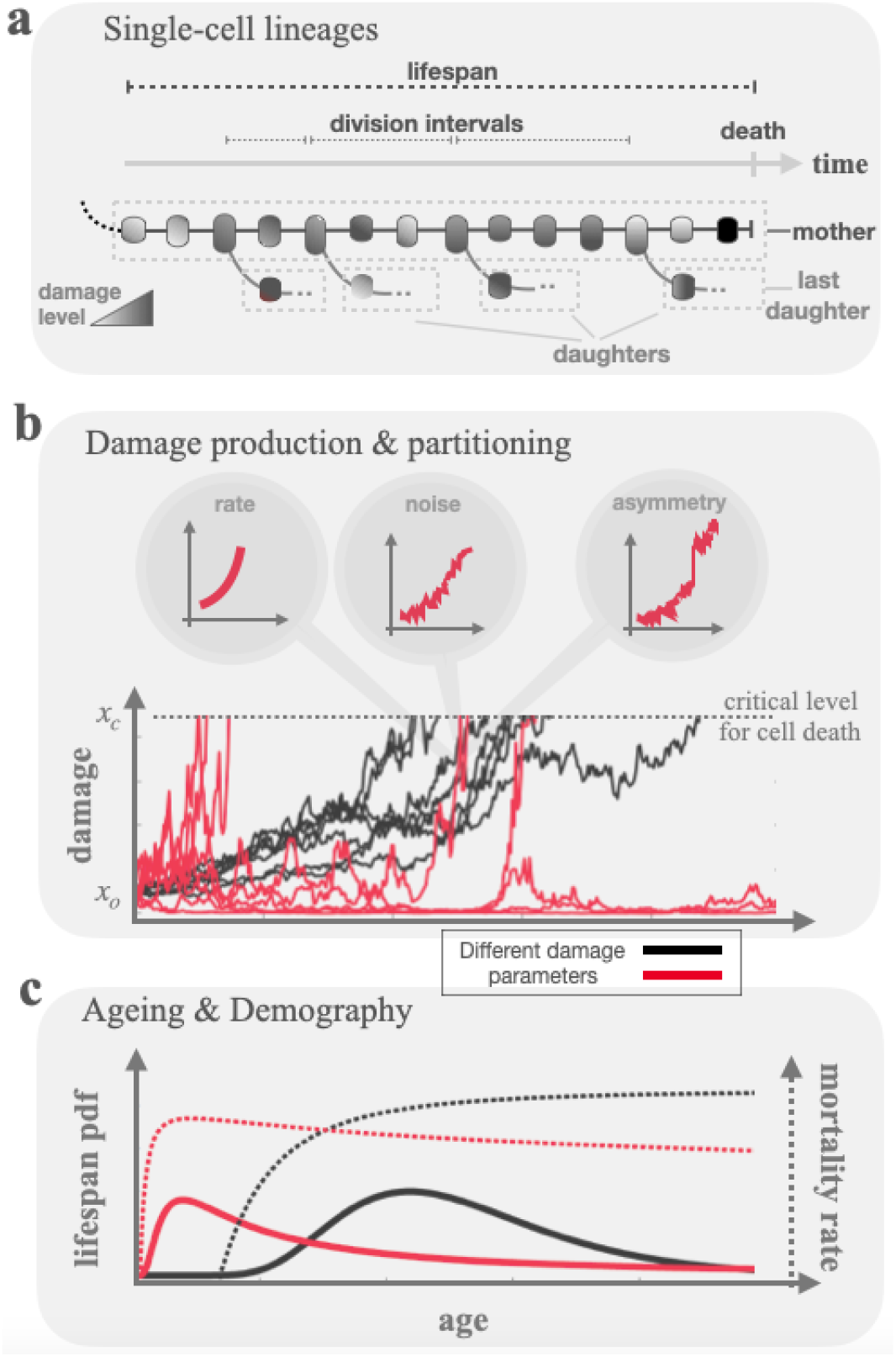
**a)** We model the damage and ageing dynamics of single-cell mother (old pole) lineages, that typically receives more damage in cell divisions. **b)** We consider a stochastic differential equation framework to model damage dynamics taking account of damage production and partition mechanisms. Each path in the figure exemplifies a damage trajectory realisation in a mother lineage where black and red colourings correspond to different cellular parameter choices. Trajectories start from the same initial damage and terminate at different times at crossing a critical damage level which defines the time of death, i.e. the lifespan. **c)** The parameters settings define the damage dynamics that in turn dictate the ageing dynamics and shape demographic patterns like the lifespan distribution and mortality rate curves.

partitioning describes the sudden jump due to asymmetric cell content (cytoplasm) partitioning during a division event. The division of timings are stochastic and drawn from a fixed distribution. Once the parameters of damage dynamics are set, initial damage levels dictate individual stochastic damage trajectories (**Fig. 1-b**). First time crossing the critical damage level determines the time of cell death from which we obtain the lifespan distribution and the mortality rate curve (**Fig. 1-c**). In this manuscript, we focus on a special case where all the parameters are constants. In other words, damage will influence the lifespan but will not affect cellular growth and division mechanisms. This choice is mostly for the sake of mathematical derivations and serves as a standard stochastic model to link cellular damage and demography. Future numerical works may relax this assumption and explore the model extensively. In the following part of this section, we present the mathematical derivations before we illustrate consequences and the applied insights of the model in the results section.

Mathematically, we consider a jump-diffusion dynamics [Merton 1976, Kou 2002, Kou and Wang 2003] which, in its most general form, is expressed by the following stochastic differential equation (SDE)

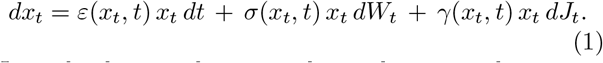

Here, the damage density *x* along a lineage realisation at time *t* is a stochastic variable and denoted as *x*_*t*_. *ε* (*x*_*t*_, *t*) is the deterministic damage production rate. *σ* (*x*_*t*_, *t*) is the strength of noise which is modelled with a Wiener (Brownian) process where *W*_*t*_ is a gaussian random variable with mean and variation as 0 and *t*, respectively. The timing of division events, and the direction and magnitude of damage jumps are modelled with a random process *J*_*t*_ and the parameter *γ*(*x*_*t*_, *t*), respectively. The *γ*(*x*_*t*_, *t*) is the sudden damage jump (asymmetry) parameter at cell division where *γ* = 0 is for symmetric division and 1 *> γ >* 0 is for a mother (old pole) cell receiving more damage than daughter cell. When a cell divides, damage density *x*_*t*_ will jump to a new damage *x*_*t*_ (1 + *γ*) along the mother cell lineage, whereas a new daughter cell will be produced with damage density *x*_*t*_ (1 *− γ*). Note that *−* 1 *< γ <* 0 corresponds to the mother’s rejuvenation, i.e. receiving less damage compared to the daughter cell. Our modelling is also valid for this scenario but will not be explored here. Cell damage level starts from an initial level *x*_0_ at time *t* = 0 and the cell mortality is defined by the first crossing of a critical (absorbing) level *x*_*c*_. Note that time and age are identical in our model therefore *t* refers to both of them.

In this manuscript, we explore the case of all parameters in the SDE as constants. Therefore, the above SDE is reduced to a simpler form:

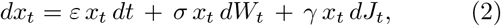

where an explicit solution is known as

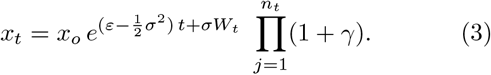

with *n*_*t*_ being a random variable for the total number of cellular divisions (jumps) until time *t* in a life trajectory realisation. The following iterative equation can be used for simulations of this jump-diffusion dynamics.

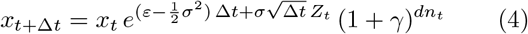

where *Z*_*t*_ is a random number from a unit normal distribution and the jump process takes *dn*_*t*_ = 1 or 0 at each discrete time step depending on the jumping process *J*_*t*_. One can simulate many individual trajectories and obtain a statistical description, but we further seek an analytical expression and rewrite the last term of the exact solution in Eq. 3 as 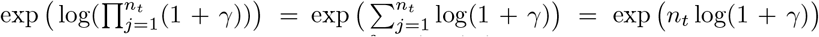, thereby transforming it into

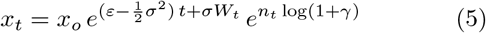

If we assume the number of divisions per unit time (i.e. division rate) follows a Gaussian distribution 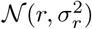, then follows also a Gaussian distribution 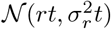. This assumption allows us to rewrite the above expression in an approximate compact solution form as

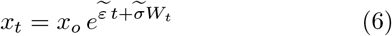

where the modified drift and diffusion terms are

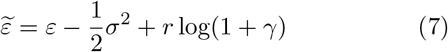

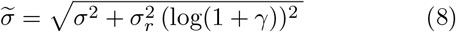

Therefore, the original SDE in Eq. 2 is approximated to the following geometric Brownian motion

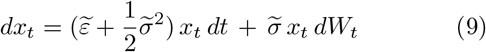

We can also express it in arithmetic Brownian form as 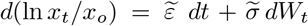, from which we obtain the probability density function for damage level *p*(*x, t*) by applying the Fokker-Planck equation with the corresponding initial and boundary conditions of *p*(*x, t* = 0) = *δ*(*x − x*_*o*_) and *p*(*x* = *x*_*c*_, *t*) = 0 as

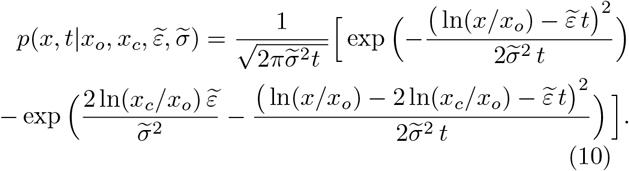

Expressing the probability density function for lifespans *f* (*t*) is equivalent to determining the first passage time to the absorbing level *x*_*c*_. Interestingly, it follows a distribution of an inverse Gaussian (IG) as

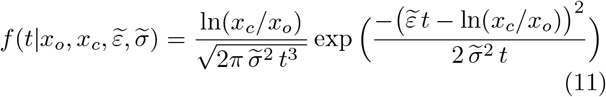

The IG nature of lifetimes was first discovered in hazard analysis by Chhikara and Folks [1977] but the importance in demography was highlighted by Anderson [2000] and Weitz and Fraser [2001], to our knowledge. Following Weitz and Fraser [2001], we rewrite this 4 parameter expression in terms of 2 parameters and make a connection from *molecular* (cellular) to *macroscopic* (demographic) parameters, i.e.

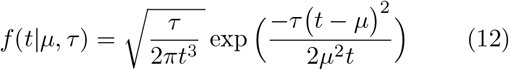

where *µ* and *τ* are respectively the mean and shape-related parameters and can be expressed in terms of cellular parameters as

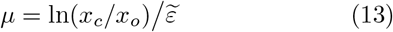

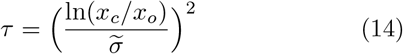

The variance of IG distribution is *µ*^3^*/τ* which is in terms of cellular parameters

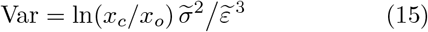

The shape of the distribution is entirely determined by *ϕ* = *τ/µ* which is

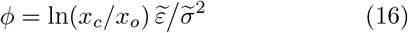

where a large *ϕ* results in a symmetric distribution. We observe that ln(*x*_*c*_*/x*_*o*_) plays a scaling factor for the mean, variance and shape of the lifespan distribution. Hence, we observe that the variance-to-mean ratio or the index of dispersion *D* of lifespans (i.e. *D* = *µ*^2^*/τ*) becomes a damage-free measure, i.e.

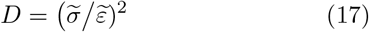

This helps interpret empirical data and molecular mechanisms as initial and critical damage levels can be difficult to quantify empirically. *D* quantifies the dispersion level of lifespans and compares it to an underlying Poisson-like mechanism where *D* = 1 (i.e. as if all death events are due to a extrinsic killing process with a constant rate). *D <* 1 and *D >* 1 indicate under- and over-dispersed distributions, respectively.

The lifespan demographic data is also represented using the function of survival probability, i.e.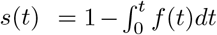. Thanks to the IG form, it can be expressed in analytical expression in our model as

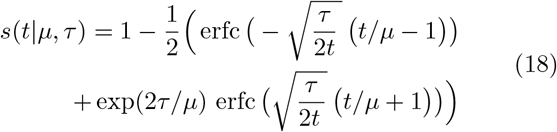

where erfc is the complementary error function. Especially the mortality rate curve is useful in an ageing context. Having obtained Eq. 12 and Eq. 18, we can calculate it using

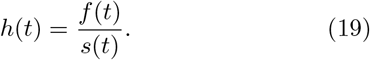

It shows how mortality chance is altering with age, where a constant mortality rate refers to a no-ageing case and an increase in mortality rates corresponds to an elevation in the underlying ageing processes. IG distributed lifespans are known to provide diverse mortality curve patterns including first increasing followed by staying constant (i.e. mortality pleautes) or by decreasing rates Weitz and Fraser [2001].

Lastly, we can also obtain a more general model where we integrate a damage-independent mortality process (similar to a Makeham term) into our modelling framework. We hold the simplest case of adding a constant killing rate *h*_*o*_ into to our mortality rate function. The modified functions for mortality rate, survival and lifespan distribution respectively become

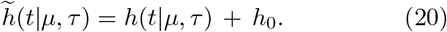

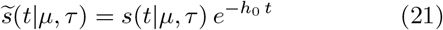

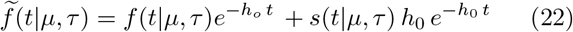

Note that the obtained mortality rate function is different than the classical phenomenological Gompertz-Makeham law of mortality which has a simple exponential form with three parameters (*α, β, λ*):

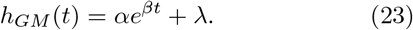

In the next section, we first explore how *micro* scale damage and cellular characteristics give rise to *macro* scale demographic patterns. Lastly, we apply our general model (the combination of the damage model with a constant killing process, Eq 22) to available empirical data.

## RESULTS

### Biologically relevant parameter ranges

Our model is relevant for single-celled systems ranging from unicellular organisms to cells within tissues. To illustrative our model’s results, we aim at biologically relevant parameter ranges of bacterial single-cells, in particular *E*.*coli*. Division rate *r* in bacteria varies depending on conditions. It reaches *r* ∼ 3.0 divisions per hour in rich media at ideal temperature whereas it is much slower under natural conditions including frequent growth arrests (*r*∼ 0). The damage partition asymmetry is controlled by parameter *γ* assuring mothers to take over more damage. We consider the full range of asymmetry levels, i.e. *γ* = 0*−* 1 where 0 refers to full symmetry and 1 full asymmetry at cell divisions. The special case of *γ* ∼ 1 can be realised but requires an active energy-dependent process [Kaeberlein 2010]. The damage production rate *ε* and fluctuation *σ* are hard to estimate as they are not directly measured. They are embedded into complex gene expression dynamics and regulatory networks that integrate transcription, translation, decay and dilution mechanisms, as well as, influenced by environment. We start with the condition that *σ* is related to *ε* with a constant noise factor. Here we consider a relative noise measure as *ξ* = *σ*^2^*/ε*, which takes values from no noise (*ξ* = 0) to high noise (*ξ* = 2). The net damage rate may range from *ε* = 0, i.e. when unwanted damage entities are efficiently eliminated or diluted, to extremely high values of *ε*, where the damage level quickly escalates resulting in a sudden mortality (e.g., a very toxic environment or under bactericidal antibiotic conditions). In order to set a numeric value for the critical damage production rate, we can refer to Eq. 13 where the case of *ε*≪ *r* log(1 + *γ*) states that the dominant damage drift factor is the asymmetric division process. Considering a typical value of a division rate *r* = 2 *−* 3 for laboratory set-up and maximal asymmetry *γ* = 1, we argue that *ε* ∼ 1 reflects high damage production. We consider *ε*≫ 1 for extreme damage rate, referring to a sudden death case which is trivial and not interesting for ageing studies, but might be of relevance for evaluating the efficacy of anti-infective substances aiming to prevent any single cells to escape treatment. We consider *ε* ∼ 0.1 for a lower damage production rate. The logarithmic ratio between the critical and initial damage level ln(*x*_*c*_*/x*_*o*_) scales time (see Eq. 13 and Eq. 15 for the mean and variance of lifespan distribution), so whenever suitable we present our results with this time scaling unit from here on.

### Analytical expressions vs numerical simulations for the dynamics of damage and ageing

In **Fig 2** we show an example of the combined dynamics of damage and ageing for particular parameter choices. The results are obtained both from our analytical expressions (Eqs. 10, 11, 19) and from numerical simulations obtained using the discrete-time version of the explicit solution for the SDE model (Eq. 4). **Fig 2-a** illustrates how cellular damage level, with its substantial along lineage stochastic fluctuations, changes over time from a defined initial value until reaching a critical value defining the time of death. We observe also substantial among lineage variance in damage levels, some trajectories following distinct patterns resulting in a variation in lifespans. The demographic patterns are shown in **Fig 2-b** with a lifespan distribution that follows an inverse Gaussian distribution and a mortality rate that exemplifies a rapid early-age increase followed by a constant probability of death at older ages. **Fig 2-c**, using both analytic and simulation methods, highlights how the demographic fates at *macro* scale are changed if the damage parameters at *micro* scales alter (for corresponding damage dynamics, see also **Fig SI-1** in SI). As pointed in the model section, they demonstrate how initial damage level scales the mean and variation of lifespan distributions, how higher damage rate and damage asymmetry shift the distribution to earlier mortalities, and how damage noise spreads the lifespans. Overall, our approximate mathematical solution is in close agreement with the numerical simulations obtained with the more explicit solution and illustrates how damage parameters influence demographic patterns.

**FIG. 2.**
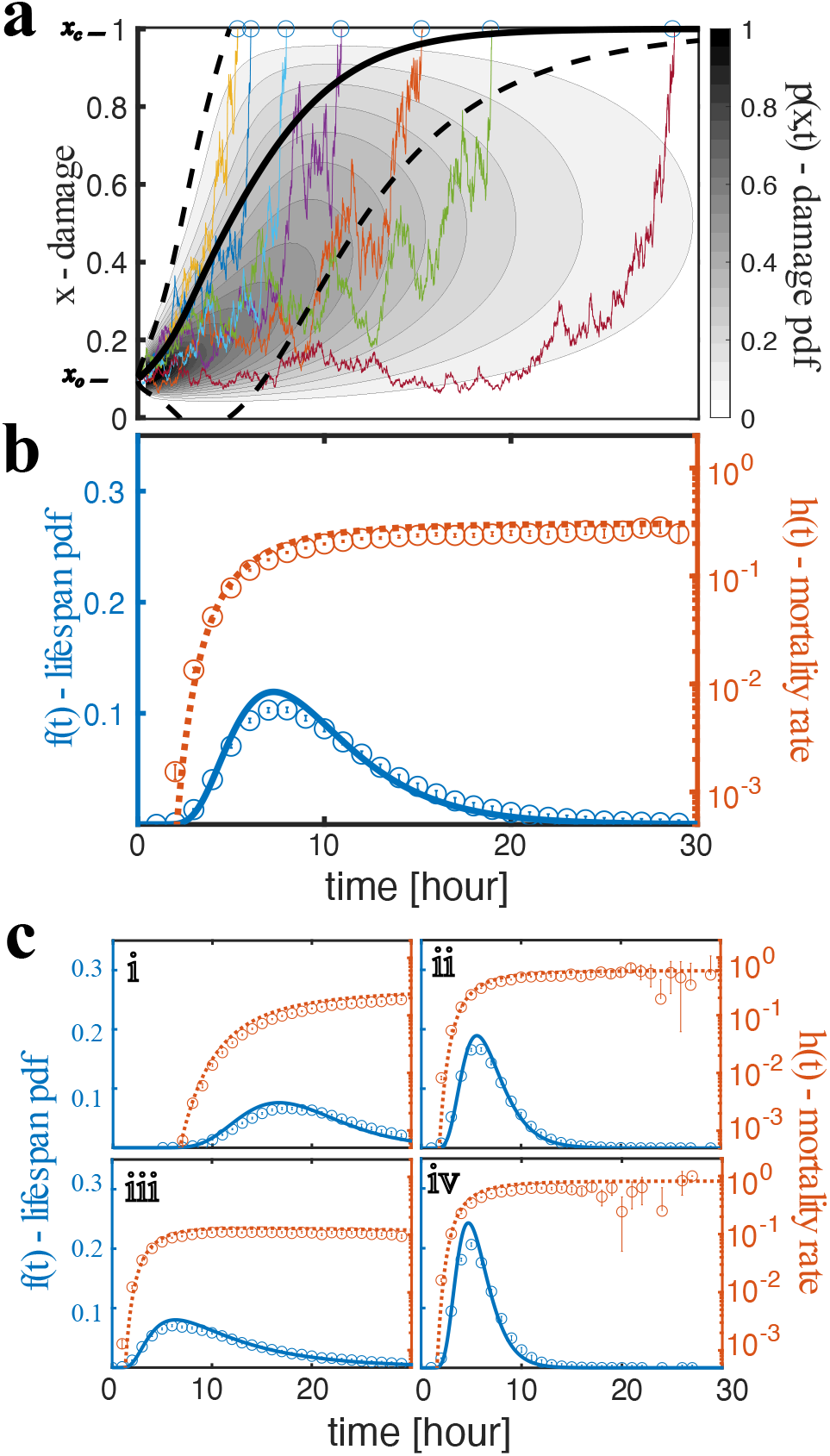
The dynamics of damage and ageing according to our model is demonstrated by using both the simulations of the exact solution (*N* = 10^5^, Δ*t* = 10^*−*2^) and the analytic expressions of the approximation method. **a, b:** Model parameters (see Table I for definitions) are *x*_*o*_ = 0.1, *x*_*c*_ = 1.0, *ε* = 0.1, *σ*^2^*/ε* = 1.0, *γ* = 0.1, *r* = 2.0, *σ*_*r*_ = 0.5. **a:** Selected damage trajectories (different colours) exemplify the stochastic realisations in the simulation. The gray contours and shading correspond to the probability distribution function of damage levels *p*(*x, t*) in Eq 10. The black solid and dashed curve stand for the mean and two standard deviation obtained from this analytic expression by numerical integration (the death cells are fixed to *x*_*c*_ for this calculation). **b:** Left y-axis: the lifespan probability density function is shown with the blue circles (simulations) and the blue curve (analytic expression, Eq. 11). Right y-axis: the same data is represented with the mortality rate measure with the orange circles (simulations) and the orange dotted curve (analytic expression, Eq. 19). **c:** This panel demonstrates how the lifespan distribution and mortality curves are affected by changing the damage parameters, e.g. **i-** initial damage decreases, i.e. *x*_*o*_ = 0.1 *→ x*_*o*_ = 0.01; **ii-** damage rate increases, i.e. *ε* = 0.1*→ ε* = 0.2; **iii-** damage noise increases, i.e. *σ*^2^*/ε* = 1.0 *→ σ*^2^*/ε* = 2.0; **iv-** damage asymmetry increases, i.e. *γ* = 0.1*→ γ* = 0.2. Error bars show 95% binomial proportion confidence intervals.

### Lifespan (mortality) statistics for different damage asymmetry and noise levels

Our analytical derivation provides an exact mapping from cellular damage parameters to demographic statistics where lifespan distribution follows an inverse gaussian function whose mean, variation, shape and dispersion levels can expressed in terms of cellular parameters (i.e. Eqs. 13, 15, 16 17). Using this mapping, we can ask how asymmetry in damage partition and stochasticity (noise) in damage production influence ageing patterns by observing the lifespan distribution and mortality rate curves. In **Fig 3** we show how these statistics change with the noise and asymmetry parameters in low and high damage rate cases. We consider a relative noise measure in damage production, i.e. *ξ* = *σ*^2^*/ε*. For the case of high damage production, both noise and asymmetry influence the lifespan distribution significantly. Higher asymmetry facilitates faster damage deposition along mother lineages and therefore shortens both the mean and variation of lifespan. Similarly increase in damage production noise will result in a drastic shift and spread of lifetimes. For the case of low damage production, the noise factor affects less, and a fast damage accumulation and early mortality is only possible if there is significant damage partitioning asymmetry, otherwise, cell lineages are likely observed as very long living, even as practically immortal if they start from initially low damage.

**FIG. 3.**
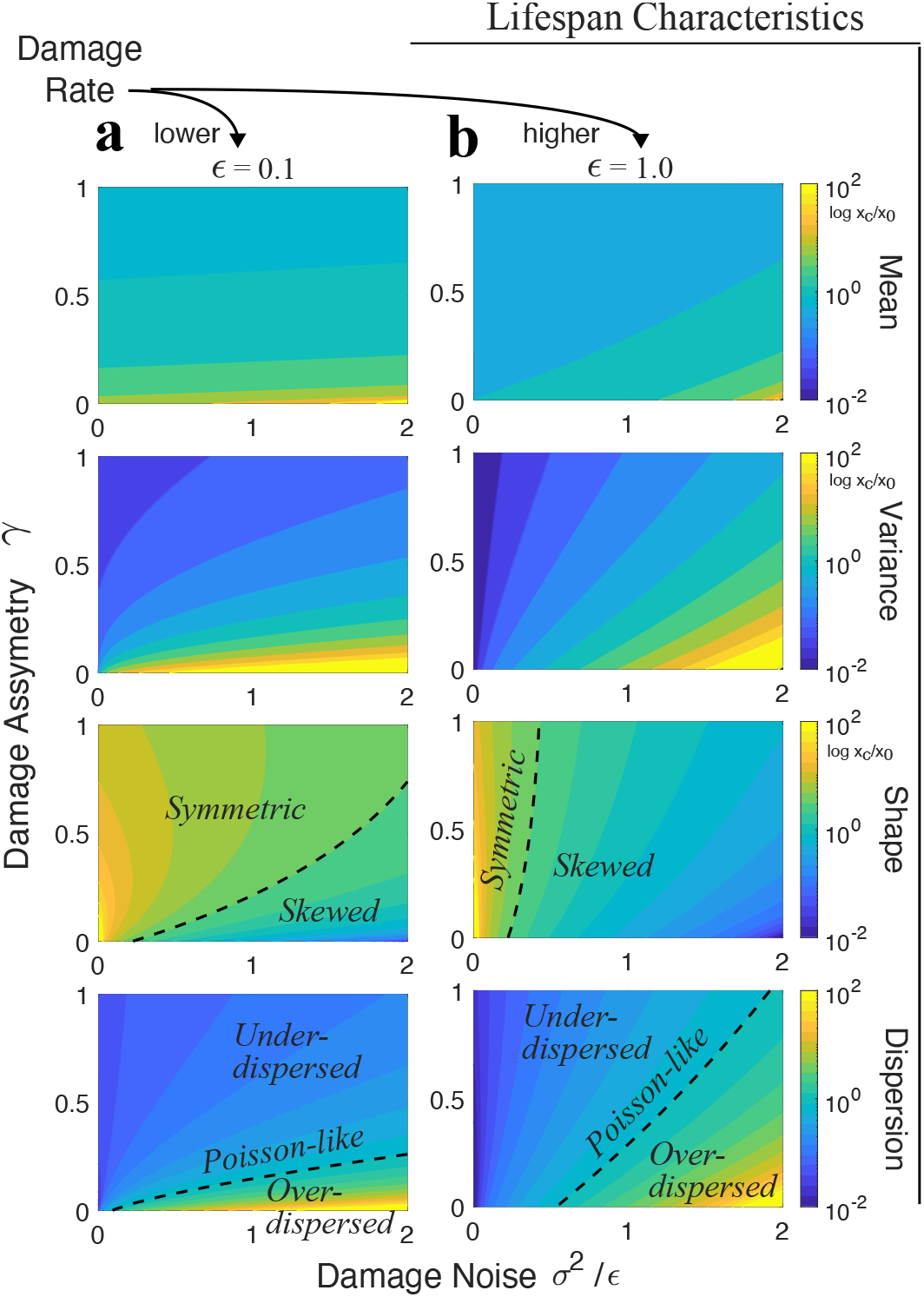
**a)** Our analytic expression maps molecular damage parameters to demographic patterns. The panels show how relative noise in damage production (measured with *σ*^2^*/ε*) and damage partition asymmetry level *γ* determine the lifespan statistics of mean, variance, shape and dispersion for a low (a: *ε* = 0.1) and a high (b: *ε* = 1.0) damage rates. Other settings for this figure are *r* = 2.0 and *σ*_*r*_ = 0.5, and we plot the results (contours) with ln *x*_*c*_*/x*_*o*_ scaling. In the dispersion plots we show the *D*∼ 1 region corresponding to Poisson-like dispersion as well as under- and over-dispersed regions (*D <* 1 and *D >* 1, respectively).

As a damage-free measure, the index of dispersion *D* (variance-to-mean ratio) of the lifespan distribution provides additional insights into the underlying mechanisms leading to demographic heterogeneity. The dispersion level of a standard Poisson process can be a reference as its *D* = 1. We can draw a direct link from molecular relative noise level (*ξ* = *σ*^2^*/ε*) to observed lifespan dispersion by adapting Eq. 17 as

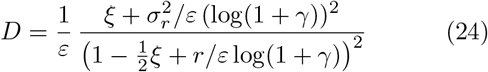

In case of complete damage partitioning symmetry (i.e. *γ* = 0), the above expression is reduced to *D ε* = (2*ξ*/ (2− *ξ*))^2^. For example, for *ε* = 1, this sets a threshold damage noise level as *ξ* = 2*/*3 which leads to a lifespan distribution as generated by a Poisson process (e.g. a constant background killing mechanism). **Fig 3** indicates which cellular parameter regions exhibit a Poisson-like behaviour (see also **Fig SI-2** in SI for flat mortality rates corresponding this *D ∼* 1 region). *D >* 1 and *D <* 1 parameter regions correspond to over- and under-dispersed lifespan distributions with respect to a Poisson process. In the under-dispersed *D <* 1 case, lifespans accumulate more closely to the population mean (e.g. as in Binomial distribution), presenting a more regular killing process with respect to randomness in Poisson process. In the over-dispersed *D >* 1 case, lifespans spread away from the population mean (e.g. as in geometric or negative Binomial distributions), presenting a more random process in comparison to a Poisson process. We especially observe that a damage production with a strong noise characteristics (*ξ* = *σ*^2^*/ε >* 1) results in a very dispersed lifespan realisations, i.e. an ever remaining chance of observing an individual with a long lifespan despite the majority of individuals might die at an earlier age.

**Fig SI-2** in SI shows the actual lifespan distributions and mortality rate curves for selected parameters. They exemplify the capability of producing different lifespan shapes including various degrees of dispersion and skewness. Furthermore, they demonstrate various mortality rate scalings with age, including the cases of a rapidly increasing scaling, an increasing scaling with a plateau, and a first increasing then decreasing scaling, where the last is not a possible pattern in classical exponential models such as in the Gompertz-Makeham model but observed in nature [Jones et al. 2014, Steiner et al. 2019, Weitz and Fraser 2001].

### Influence of division rate distribution on lifespan statistics

Our model provides also a direct inverse relationship between division rate and lifespan (Eq. 13). Keeping all other parameters fixed and having an asymmetric partition case (*γ >* 0), a higher division rate will result in higher damage accumulation in mother lineages causing earlier death events, i.e. shorter lifespans. The inverse relationship between mean lifespan and mean division rate is demonstrated in **Fig 4-a**. This observation is in agreement with the classical pattern observed between reproduction and longevity [Kirkwood and Rose 1991]. However, reducing mean division rate is likely not a permanent escape from mortality in an evolutionary strategy as it trades off with the fitness components such as lifetime reproductive output, i.e. the number of offspring (division events) over the entire lifespan. In general, the fitness consequences for population growth can be more complex which requires further exploration. Nevertheless, we highlight how the heterogeneity of lifespans depends on the mean and variation of division rate in our model. We consider here the dispersion level of the lifespan distribution as a measure of heterogeneity in lifespans as computed with Eq. 24. **Fig 4-b** demonstrates this relation and shows us that a decrease in mean division rate as well as an increase in the variation of division rate drives lifespan dispersion. This can be interpreted as that cells growing in optimal conditions (e.g. exponential phase) probably yields in higher division rate metabolism and keep an homogenous lifespan distribution in a population. This uniform demographic fate is perturbed when the cells’s division rate is reduced or become noisier, presumably due to non-optimal or limited conditions (e.g. stationary phase or under extrinsic stress). At these limits, not only the mean of lifespan is shifted but also more heterogeneous group of individuals are expected to be observed which can be beneficial for the survival of a population.

**FIG. 4.**
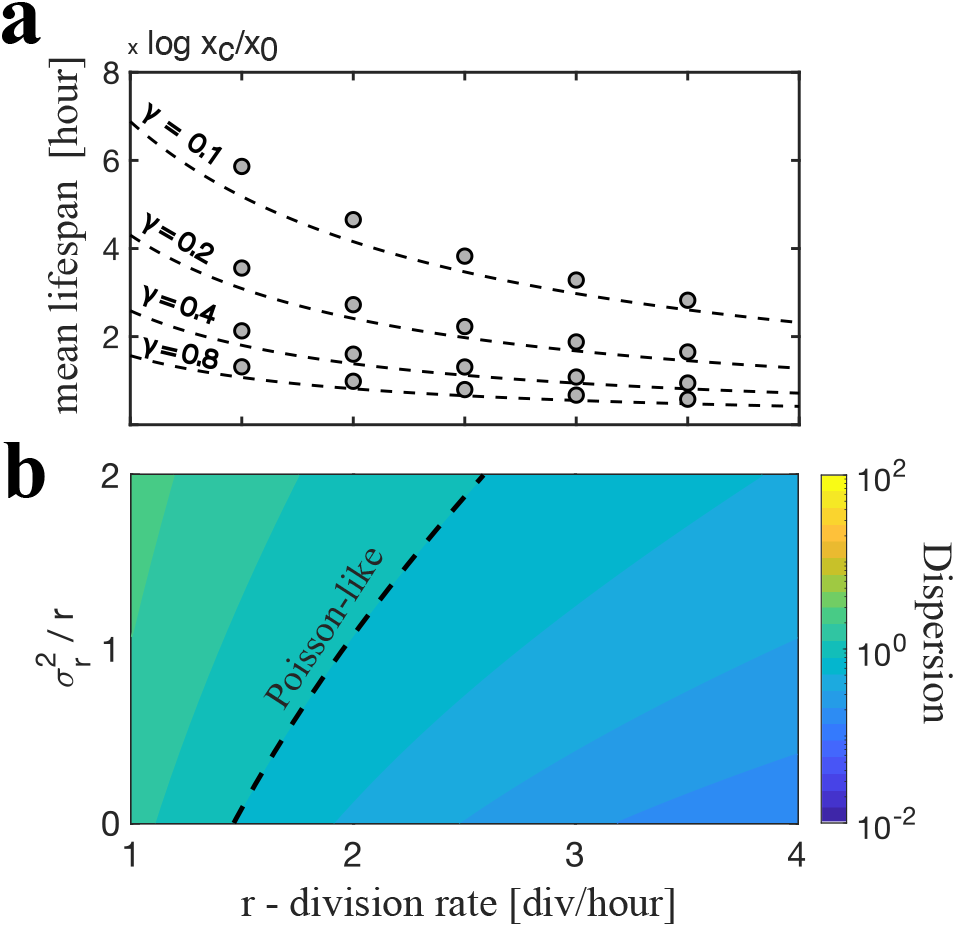
**a-** Mean lifespan is inversely proportional to the mean division rate *r* when an asymmetry exists in damage partitioning. Scaling with different asymmetries *γ* = 0.1, 0.2, 0.4, 0.8 are shown for analytical expression (dashed curves) and for simulation averages (circles, 95% CI as error bars) for parameters are *x*_*o*_ = 0.01, *x*_*c*_ = 1.0, *ε* = 0.1, *σ*^2^*/ε* = 1 and 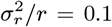. **b-** Lifespan dispersion *D* depends on both the mean and the variance of division rate. We use the same parameters as in **a** and fix *γ* = 0.2. The dashed line is for *D* = 1 corresponding to Poisson-like dispersion level.

### Application to empirical data

Finally, we revisit four previously published datasets on single-cell *E*.*coli* ageing, obtaining their empirical mortality statistics and employing our stochastic damage model (combined with a constant killing factor) in comparison with the Gompertz-Makeham model. These studies compared the longevity of mother cells of a wild type *E*.*coli* strain with the mother cells of a mutant case (lexA3 for Wang *et al*. [2010]; mutH strain for Robert *et al*. [2018] and rpos deletion for Yang *et al*. [2019]), or with the last daughter cell of a mother cell lineage in Steiner *et al*. [2019]. All these interventions provide us with essential quantitative data to make general observations on ageing characteristics, in particular on mortality scalings with age (or lifespan distributions).

To obtain empirical survival scaling with age, we used the Kaplan-Meier (K-M) estimation method and incorporated right censorship. To ensure a fair comparability across studies, before we apply the K-M estimate, we binned the lifespan data in 6 hour intervals, as guided by the Freedman-Diaconis rule. We fit our damage model (using the modified version including a constant background killing rate, i.e. Eq. 22) by a maximum likelihood method on the actual (not binned) lifespan and censorship data. Similarly, we also fit the Gompertz-Makeham model (Eq. 23) for comparison. **Fig. 5** shows both the empirical and model fitting results for the mortality rate scaling with age, as this measure acknowledges ageing characteristics more clearly (see also **Fig. SI-3** in SI where the same data are presented as lifespan distribution). Our model fits well the empirical scalings of various datasets. For most of the cases (7*/*8), it becomes the better supported model against the classical Gompertz-Makeham model, based on the Akaike information criteria (AIC) (see Table II).

**TABLE I.** Key Notations.

| Notation | Description |
| --- | --- |
| $x_t$ | Stochastic damage density at time $t$ |
| $x_o$ | Initial damage |
| $x_c$ | Critical damage |
| $\varepsilon$ | Damage production rate |
| $\sigma$ | Noise strength in damage production |
| $\gamma$ | Asymmetry level (0 = Symmetry) |
| $r$ | Mean division rate |
| $\sigma_r$ | Standard deviation of division rate |
| $W_t$ | Wiener (Brownian) process |
| $J_t$ | Jump process |
| $s$ | Survival probability |
| $f$ | Lifespan probability density function |
| $h$ | Mortality rate function |
| $D$ | Index of dispersion of lifespan distribution |

**TABLE II.**
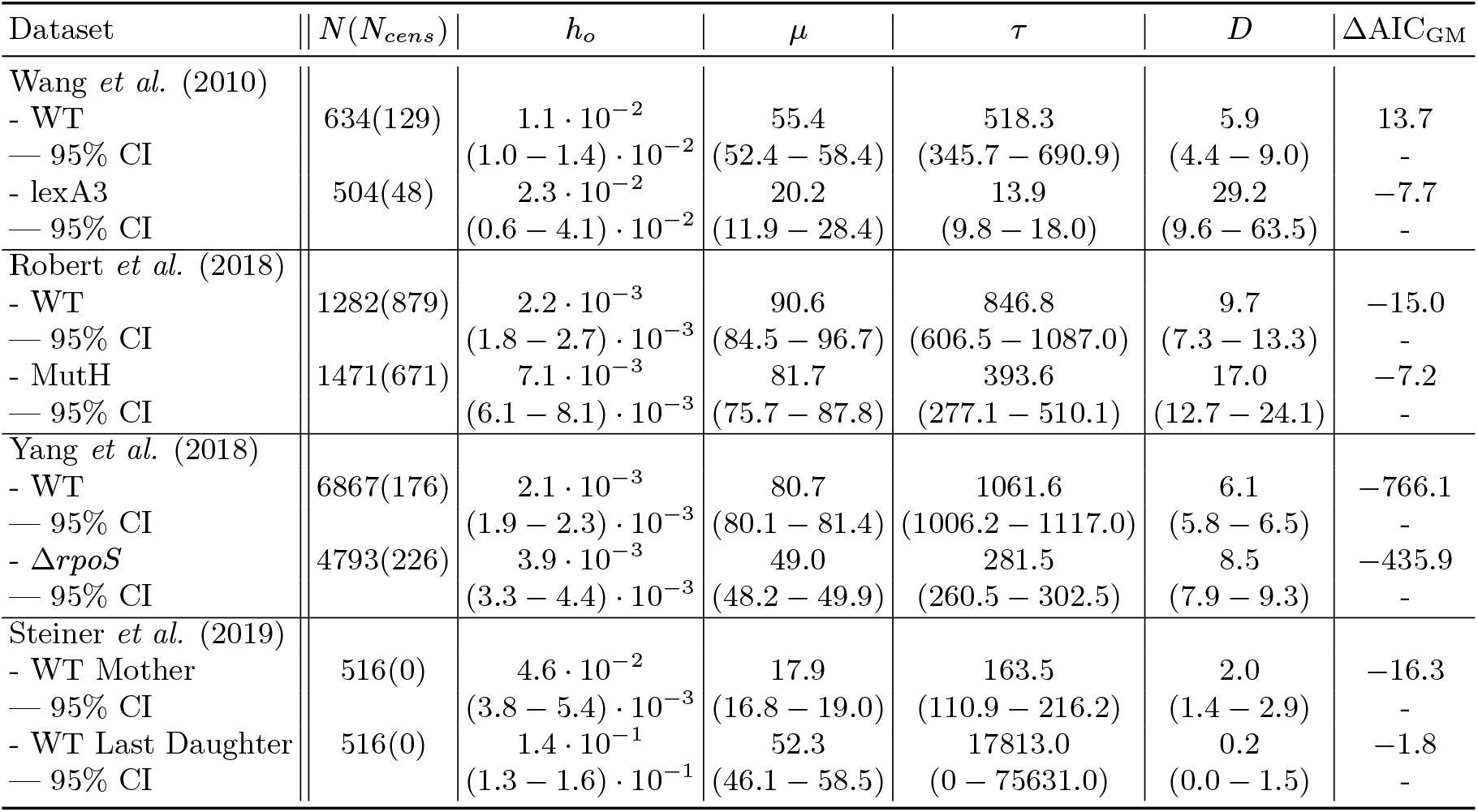
Maximum Likelihood Estimations of the Model.

**FIG. 5.**
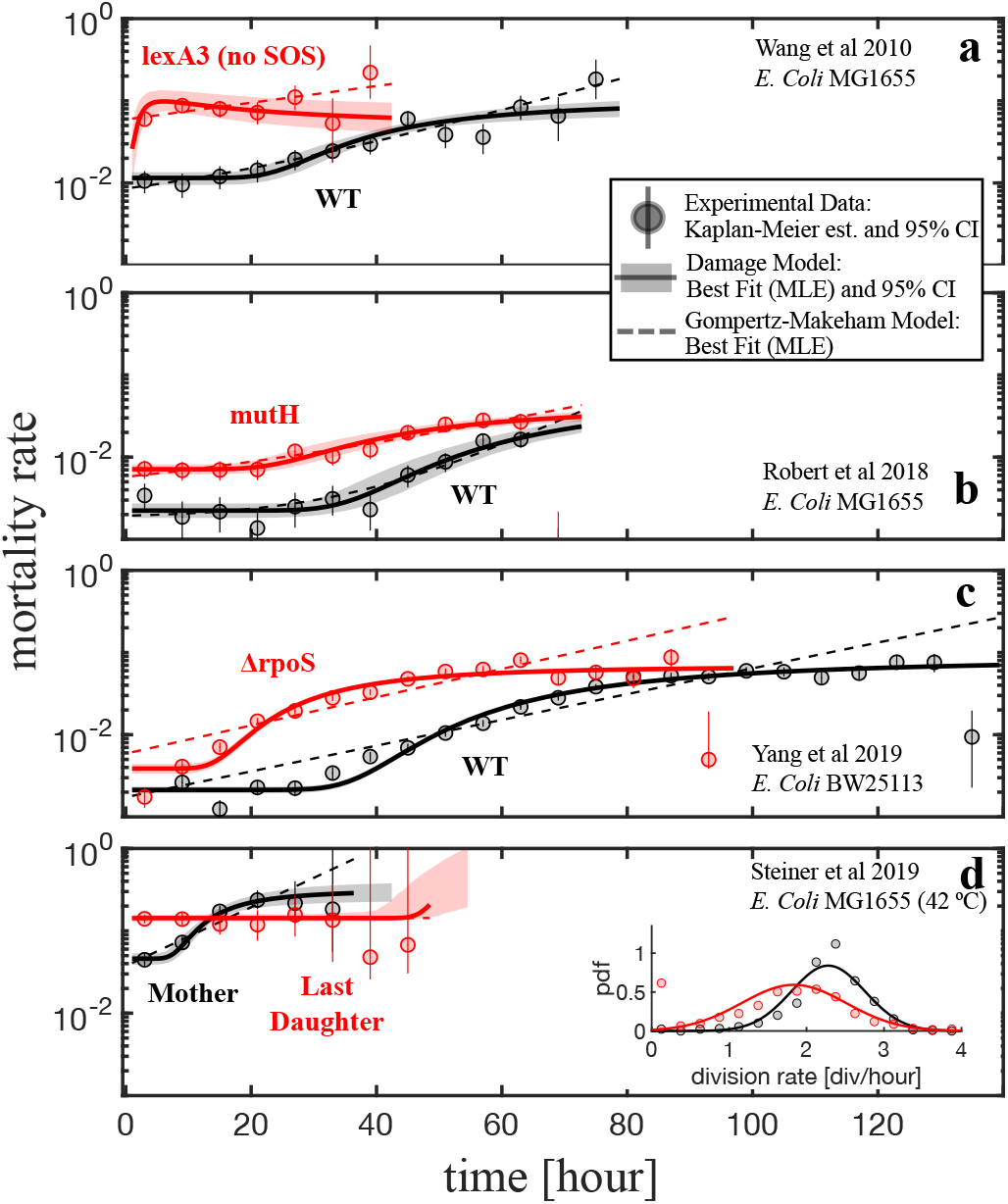
Mortality rate data for *E*.*coli* single-cell data from four different studies: **a)** Wang *etal*. [2010]. **b)** Robert *etal*. [2018] **c)** Yang *etal*. [2019] **d)** Steiner *etal*. [2019]. The Kaplan-Meier (K-M) method was used to estimate the empirical rates (circles) with their 95% CI (error bars) after binning the data in 6 hour intervals (the Freedman-Diaconis rule). Our damage model with a constant external killing rate (Eq. 22, solid curves with shading for 95% CI) and the Gompartz-Makeham model (Eq. 23, dashed curve) was fitted to the actual (not binned) data using a maximum likelihood method. **Inset:** The division rate distributions among different lineages (bin= 1*/*4 hour) and the fitted gaussian pdfs are shown for the mother (black) and last daughter (red).

The mortality rate curves exhibit a clear profile of a late-age plateau after an early-age exponential increase for the wild strains, as demonstrated in our damage model and previously discussed [Weitz and Fraser 2001]. The basal mortality rate, which corresponds to a damage-independent killing rate in our model, and the critical age, which separates the higher and basal mortality rates, alter for each dataset. These differences can be due to differences in experimental settings and in the resulting damage mechanisms. Steiner *et al*. [2019] dataset especially shows how environmental factors such as high temperatures can accelerate the ageing dynamics, probably by increasing overall metabolic rates that leads to higher damage production rate. The critical age for the wild type in the other datasets (i.e. at standard conditions) ranges in 25 *−* 40 hours. Before this range, i.e. in the first 24 hours, mother cell lineages show a non-ageing profile, i.e. mortality rate is close to a constant basal value. This might suggest that typical damage production rate is low in standard conditions for *E. coli* where ageing kick off only at a later time point, unless a stressful environment or a mutation exists. The comparison cases for each datasets show that an intervention typically both shifts the critical age to earlier times and increase the basal killing rates. In parallel, these interventions increase the dispersion or heterogeneity of lifespans attributed to damage dynamics (measured with the index of dispersion *D* in Table II) at these interventions. Recalling that our model’s *D* is a damage-free measure (Eq.17), these estimations indicate that the initial damage load difference alone cannot explain the observed differences between comparison groups. This suggests that interventions likely induce the rate and noise in cellular damage production or change the damage partitioning characteristics with asymmetry or division mechanisms. It is striking that the reported division rate distributions in Steiner *et al*. [2019] show clear difference between mother and last daughter lineages as shown in **Fig. 5-d-Inset**. The mean division rate is reduced in the last daughter (*r* = 1.8 div/hour in comparison to *r* = 2.3 div/hour of the mother lineages) and the variation in division rates are substantially increased (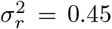 in comparison to 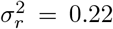 of the mother lineages). Although this quantitative justification is not a rigorous due to many unknown damage parameters, our model is instructive to guide the interpretation of the empirical data.

## DISCUSSION

Single-cell ageing studies produced remarkable quantitative data in recent years. Despite this exquisite data we need complementary mathematical models to draw the causative link between cellular damage and ageing. In particular, we need a better understanding of how damage characteristics affect demographic patterns. Such linking requires bridging noisy molecular level process to individual heterogeneity at a macro scale, even when genetic and environmental factors are fixed. To fill this gap, we introduced a generic damage model using stochastic differential equations (SDEs) accounting for damage production rate and noise strength as well as damage partition asymmetry along mother cell lineages. We focused on a constant parameter case of the SDE (Eq. 2) and successfully obtained interpretable analytical solutions for the damage dynamics (Eq. 10) and the demographic lifespan distribution (Eq. 12). Our analytical expressions match numerical simulations (as exemplified in **Fig 2**) and prove to be instructive in understanding of how *molecular* damage dynamics scale up to *macroscopic* demographic patterns. Our modelling with jump-diffusion SDE results in the lifespan distribution having an inverse-Gaussian (IG) form (Eq.12) where its statistics can be expressed explicitly with cellular damage parameters as well as inferred from empirical lifetime data using standard maximum likelihood method. Such models directly connect molecular damage parameters to demographic measures, providing mechanistic estimates in cellular processes. IG lifespan distribution and its potential application in demographic data were discussed by Anderson [2000] and Weitz and Fraser [2001] with their arithmetic Brownian SDE modelling of vitalities for classical demographic data. Our modelling extended their approach to single-cell ageing by incorporating damage partition asymmetry. As highlighted in **Fig 3**, we showed how asymmetry and relative noise affect the lifespan distribution where particular arrangements for these parameters can yield in different lifespan characteristics. This not only determines the early and late life mortality expectations but also shapes the distribution with respect to a standard Poisson-like process (as similar, or over- and under-dispersed lifespans). Such diverse lifespan behaviours alter the population and evolutionary dynamics, and provide expectations of the maintenance and evolution of heterogeneity. Apart from the mathematical contributions we developed, our modelling can be insightful for quantitative understanding in immortality conditions of bacterial cells [Wang *et al*. 2010, Rang *et al*. 2011, 2012, Łapińska *et al*. 2019]. We quantitatively exemplified in **Fig 3** that if the bacterial cells start from damage-free initial levels and are kept under low damage production conditions, one can observe very large lifespans (i.e. practically immortality) of a mother lineage unless a very extreme asymmetry exists. Typical demographic models such as the Gompartz-Makeham model are largely phenomenological and do not provide mechanistic interpretations. They are limited to exponential mortality rates and miss for example a mortality plateau pattern as observed in the bacterial lifespan data which is captured by our model as revealed in **Fig 5**. In this empirical data application, we also revealed (**Fig 5-d-inset**) how different variation levels in division rates can exist in mother and last daughter cell lineages which exhibit different lifespan characteristics [Steiner *et al*. 2019]. This might indicate a different transmission of heterostasis to offspring born to older individuals, probably triggering the noise in division decisions in cells.

Our approach directly models the stochastic dynamics of cellular damage and is not bound to an equilibrium assumption as largely held in previous bacterial ageing modellings [Watve *et al*. 2006, Evans and Steinsaltz 2007, Chao 2010, Blitvić and Fernandez 2020]. It is therefore promising to explore phenomena responsible for individual (cell-to-cell) heterogeneity observed in ageing. Studies with single-cells show rich intrinsic cell-to-cell variation in gene expression [Elowitz *et al*. 2002, Foreman and Wollman 2020, Urchueguía *et al*. 2021]. One application of our model is to relate damage to gene regulatory networks to predict cell-to-cell variation in gene expression. The relation between gene expression and ageing has been investigated mostly based on transcriptome analyses on humans and other model organisms. The importance of down-regulation of genes encoding mitochondrial proteins; downregulation of the protein synthesis machinery; dysregulation of immune system genes; reduced growth factor signalling; constitutive responses to stress and DNA damage; dysregulation of gene expression and mRNA processing, have been already underlined [Frenk and Houseley 2018]. However, to our knowledge, no connection between stochastic dynamics of gene expression and individual heterogeneity in ageing has been investigated. Our modelling can also be insightful for designing experiments to better understand the impact of cellular damage. Simultaneous measurements of both cell fates and damage-related gene expression signals are promising [Sampaio et al. 2022] where we can utilise the stochastic features of gene regulation to grasp underlying mechanisms and to inform on our modelling choices.

The analyses in this manuscript were limited to a constant parameter case in the generic stochastic differential equation setting. Future applications can relax this assumption and explore, at least with numerical simulations, other plausible biological scenarios. Time or damage-dependent parameters can be integrated into our model as damage production rate, noise levels, and asymmetry levels, and division rate might alter with cellular damage levels. Such extension will not only make our modelling framework more comparable to existing models by others [Chao 2010, Blitvić and Fernandez 2020] but also will provide elaborative results on evolutionary trade-offs between reproduction and longevity, as extensively discussed in evolutionary theories of ageing [Kirkwood and Rose 1991]. Our assumption on a critical damage level defining the time of cell death can also be altered in future applications, for example, by using a killing rate depending on the damage level such as in the form of a linear function [Evans and Steinsaltz 2007], or of a Hill function as typically observed in cellular environments due to the thermodynamics of finite particles. A more direct extension of our modelling and analyses would be to project damage dynamics into population dynamics. This will require following the damage dynamics of an entire population descending from a single *mother* cell where each newborn daughter is itself a mother lineage starting from a different initial damage level. Albeit challenging in rigorous calculations, approximations may bring us valuable grounds for answering population-related and evolutionary questions such as damage distribution in population or the optimal damage characteristics for optimized fitness [Watve et al. 2006, Evans and Steinsaltz 2007]. This theoretical achievement will complement the recent attempts to collect data simultaneously at single-cell tracking and batch population [Bakshi et al. 2021] and will advance our evolutionary understanding ageing and beyond.

## ACKNOWLEDGEMENTS

We thank Mathias Franz, Audrey M. Proenca, Shripad Tulkapurkar and David Steinsaltz for comments on a previous draft of the manuscript. US was funded by the Deutsche Forschungsgemeinschaft (DFG, German Research Foundation) – 430170797 as a Heisenberg Fellow. MT was funded by the Deutsche Forschungsgemeinschaft (DFG, German Research Foundation) – 430174701 and by the European Union under the Marie Skłodowska-Curie Actions’ European Postdoctoral Fellowship grant agreement – 101069035.

## SUPPORTING INFORMATION

**FIG. SI-1.**
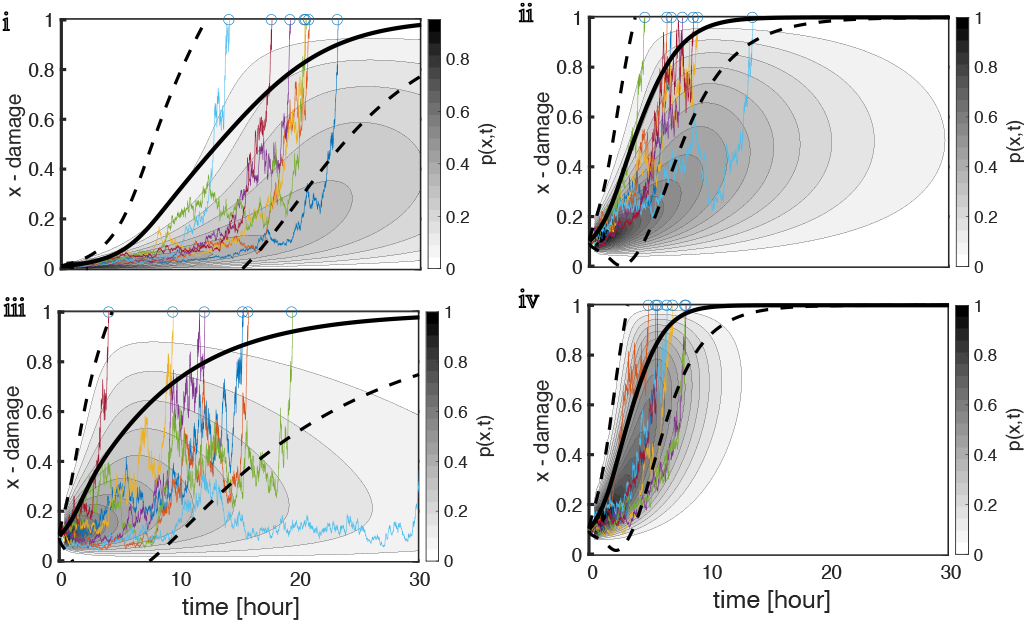
Damage dynamics is plotted for the content of **Fig** 2’s panel c. See the caption therein for the details.

**FIG. SI-2.**
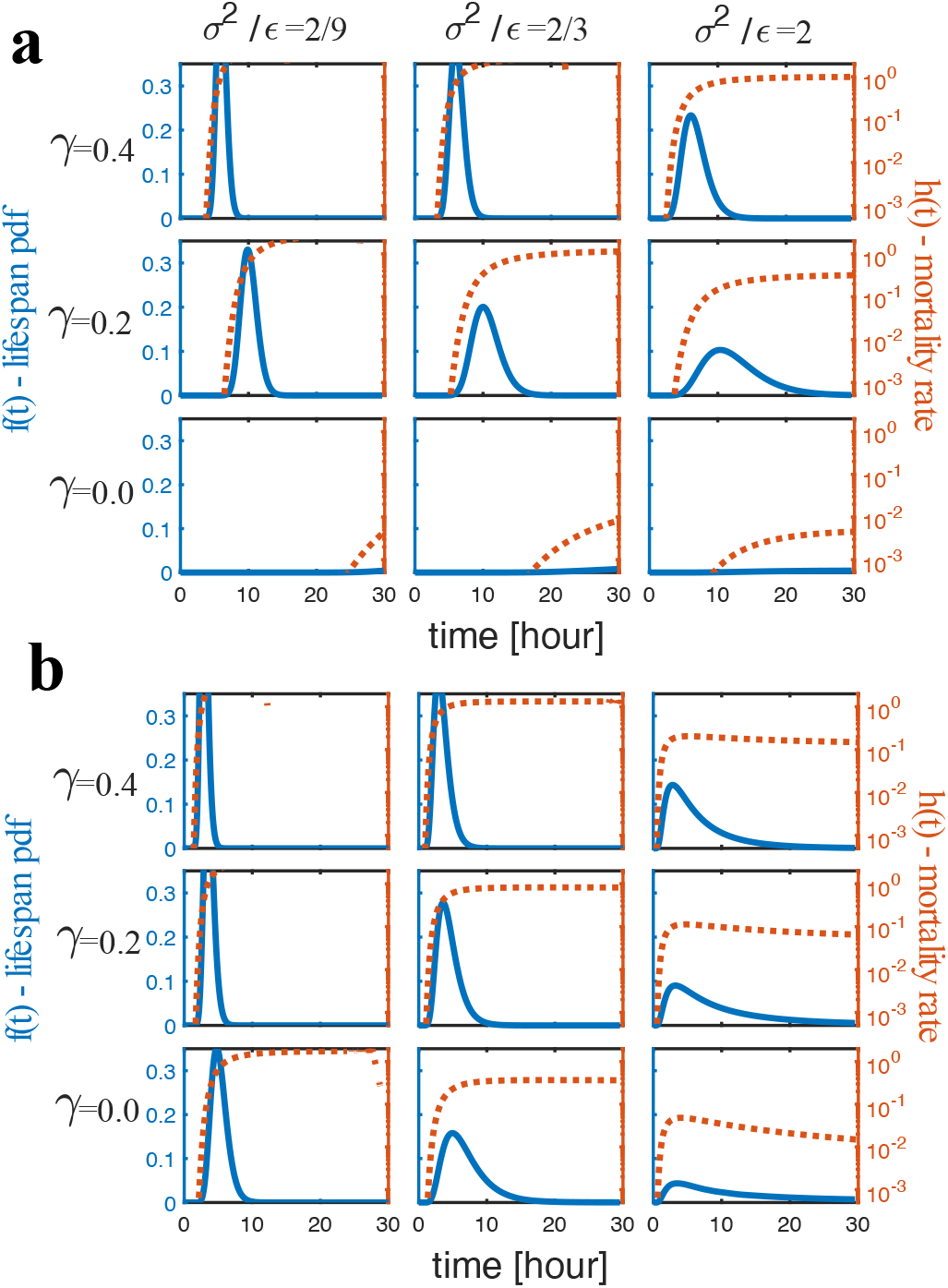
Example lifespan distributions and mortality curves are plotted for the content of **Fig** 3. See the caption therein for the details. Model parameters (see Table I for definitions) are *x*_*o*_ = 0.01, *x*_*c*_ = 1.0, *r* = 2.0, *σ*_*r*_ = 0.5. For **a:** *ε* = 0.1 and **b:** *ε* = 1.0

**FIG. SI-3.**
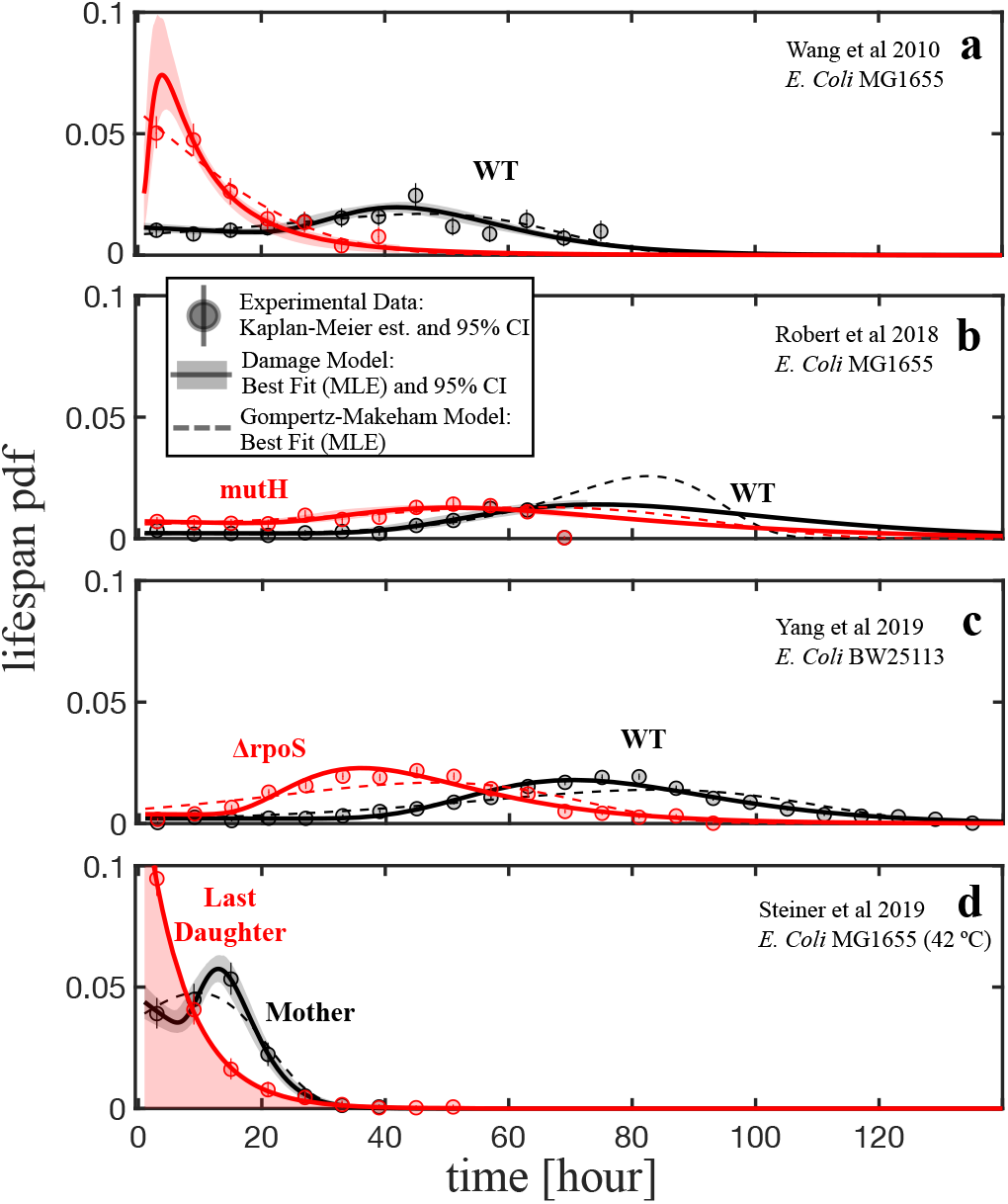
**Fig** 5’s content is replotted here in terms of lifespan pdf. See the caption therein for the details.

